# Doubling the Field of View in Common-Path Digital Holographic Microscopy via Wavelength Scanning and Polarization Gratings

**DOI:** 10.64898/2026.04.03.716314

**Authors:** Aleksandra Piekarska, Mikołaj Rogalski, Marzena Stefaniuk, Maciej Trusiak, Piotr Zdańkowski

**Affiliations:** Institute of Micromechanics and Photonics, Warsaw University of Technology, Warsaw, 02-525, Poland; Laboratory of Neurobiology, BRAINCITY, Nencki Institute of Experimental Biology of Polish Academy of Sciences, Warsaw, 02-093, Poland

**Keywords:** shearing interferometry, grating interferometry, common-path interferometry, quantitative phase imaging, phase retrieval, replica phase removal, digital holographic microscopy

## Abstract

Digital holographic microscopy systems in a common-path configuration, compared to systems with a separate reference arm, offer a compact design and resistance to disturbances. They can operate with partially coherent illumination, reducing speckle noise. However, they are limited by the overlapping of the object beam and its laterally shifted replica. As a result, images from different regions of the object overlap on the detector, preventing imaging of dense samples. We present the wavelength-scanning replica-removal method, which solves this problem by enabling the separation of information from both replicas and thereby doubling the effective field of view (FOV). The wavelength-scanning multi-shear replica removal algorithm plays a key role in reconstructing the undisturbed phase from a series of holograms recorded with variable shears. The shear value is controlled by changing the illumination wavelength. This enabled the development of two measurement modes: time-domain wavelength scanning for high-quality imaging, and a single-shot mode with frame division into color channels to improve temporal resolution. The method was validated using resolution tests and biological samples - neurons and dynamic yeast cultures. By combining the advantages of the common-path configuration with dense-structure imaging and dynamic processes, the proposed method constitutes a versatile tool for quantitative phase microscopy.

## Introduction

Quantitative phase imaging (QPI) is a label-free imaging method for the investigation of transparent cells and tissues^1^. By measuring the delay of light caused by the semi-transparent sample, QPI enables quantitative reconstruction of sample parameters such as the refractive index and dry mass^2^. This method is widely applied in the analysis of cell morphology^3,4^ and the monitoring of physiological processes^5,6^, which is of key importance for the evaluation of drug efficacy^7,8^ and the development of new therapeutic strategies^9,10^.

One of the most widely used QPI technique is digital holographic microscopy (DHM), which achieves phase measurement by decoding the optical fringe patterns, that are produced via the interference of at least two optical beams. In its standard implementation, the interfering object and reference beams propagate along separate paths^2^. This configuration has significant drawbacks: large size and susceptibility to environmental disturbances, such as temperature fluctuations or mechanical vibrations. Additionally, due to substantial differences in optical path lengths across the entire field of view, maintaining consistently high contrast requires the use of highly coherent light sources. This, in turn, leads to undesirable phenomena such as speckle noise and reduced image resolution^11,12^.

An alternative to classical, reference beam DHM, is a common-path configuration^13–20^, in particular shearing interferometry^15,16^, in which the object beam interferes with its spatially shifted copy. This ensures nearly identical optical paths, which eliminates the separate reference arm and simplifies the setup. Moreover, it enables operation under reduced-coherence illumination, while maintaining high interferogram contrast. The lateral shift is implemented, among others, using shear plates^21–27^, diffraction gratings^28–33^, Wollaston prisms^34^, or beam displacers^35,36^.

Reconstruction-wise, common-path DHM systems can be divided into two groups, depending on the magnitude of the introduced shear. The first comprises gradient-based methods, in which the shear value is smaller than the lateral size of the measured objects - typically close to the spatial resolution of the system.

In this configuration, the interferogram encodes the spatial derivative of the object phase; Therefore, recovering the full phase information requires recording the gradient components in two directions and an additional numerical integration procedure. To this end, stringent conditions must be satisfied: orthogonality of the shear vectors^37,38^ and precise knowledge of their values^39^ - identical for both vectors. Simple solutions employed for this purpose are susceptible to noise accumulation and error propagation^40–42^, whereas advanced algorithms are time-consuming and often require multiple holograms^39^. The recently proposed optimization-based approach relaxes the requirement of orthogonality of the shear vectors and precise knowledge of their values. However, it still requires a good initial estimate of these parameters, is more computationally expensive, and requires hyperparameter fine-tuning^43^.

The second group of methods are the total shear configurations^17,34,35,44–46^, in which the lateral displacement between interfering beams is larger than the lateral dimensions of measured objects. For this configuration it is extremely important that measured area is either only sparsely covered with the measured objects, or it contains a large object-free area. Otherwise, images from different regions of the sample will overlap on the detector with the laterally shifted replica, causing mutual signal annihilation and preventing the imaging of dense samples. For this reason, samples for which the size of the required reference region may be insufficient cannot be correctly imaged using this method.

To extend the applicability of total shear methods to dense and extended samples, there were proposed methods that rely on modifying one of the interfering beams to remove object information^18,20^. Although they enable imaging of dense and complex objects, they result in increased optical system complexity^47^ and a significant reduction of the effective field of view, often by more than half ^30–32^. In turn, the use of low-pass filtering by means of a pinhole for this purpose renders the system difficult to precisely align and sensitive to mechanical instabilities^18,20,48,49^.

Another solutions for imaging complex samples consists of employing shear scanning to separate overlapping fields. These include, among others, Cepstrum-based interference microscopy (CIM)^50–52^and R2D-QPI (replica-removed doubled-FOV QPI)^53^. The CIM approach relies on highly precise orthogonal single-pixel shears for temporal scanning, and the high-pass filtering, which increases system complexity and might suppress low-frequency phase components. Contrary, the R2D-QPI approach, recently proposed by our group, constitutes a general methodology that enables separation of overlapped phase information from a set of at least 2 phase images with arbitrarily chosen shears via the proposed multi-shear replica removal (MRR) algorithm. This methodology, can be implemented in various total shear hardware configurations, providing they can generate interference patterns with variable shear values.

The current implementation of R2D-QPI method was based on polarization grating total shear configuration, where the variable shear was achieved via axial scanning of one of the gratings. However, this introduced significant limitations to the system: mechanical wear, vibrations, and limited acquisition speed. The requirement to record a sequence of frames precludes the possibility of imaging dynamic samples.

In this work, we propose the wsR2D-QPI (wavelength scanning replica-removed doubled-FOV QPI) method. In this approach, controlled variation of the replica separation is achieved by changing the illumination wavelength, completely eliminating the mechanical movement within the system. At the same time, the reconstruction algorithm has been modified to compensate for the resulting chromatic aberration. The solution preserves all the advantages of the common-path configuration and the MRR algorithm (field-of-view doubling, imaging of dense samples). At the same time, it offers significant improvements: increased stability and repeatability, simplified architecture, the possibility of noise averaging, and, in an implementation with a color camera, a reduction in the required number of frames to a single frame per reconstruction. This translates into a substantial increase in temporal resolution, enabling the observation of dynamic processes.

## Results

### 2.1 Experimental Setup and Reconstruction Strategy

The common-path configuration described in this article (Fig. 1(a)) is based on the use of the polarization grating (PG) module^17,43,46,53^, built from two identical polarization gratings arranged in series. It splits the illuminating beam into two diffraction orders (±1) with orthogonal circular polarizations and generates a lateral wavefront shear τ_*λ*_. Rotation of one grating around optical axis changes the deflection angle of the beams at PG2, adjusting the beam inclination and the spatial frequency of the interference fringes. In the measurements, we set a standard carrier frequency for the phase reconstruction using the Fourier transform method (FTM).

**Fig. 1.**
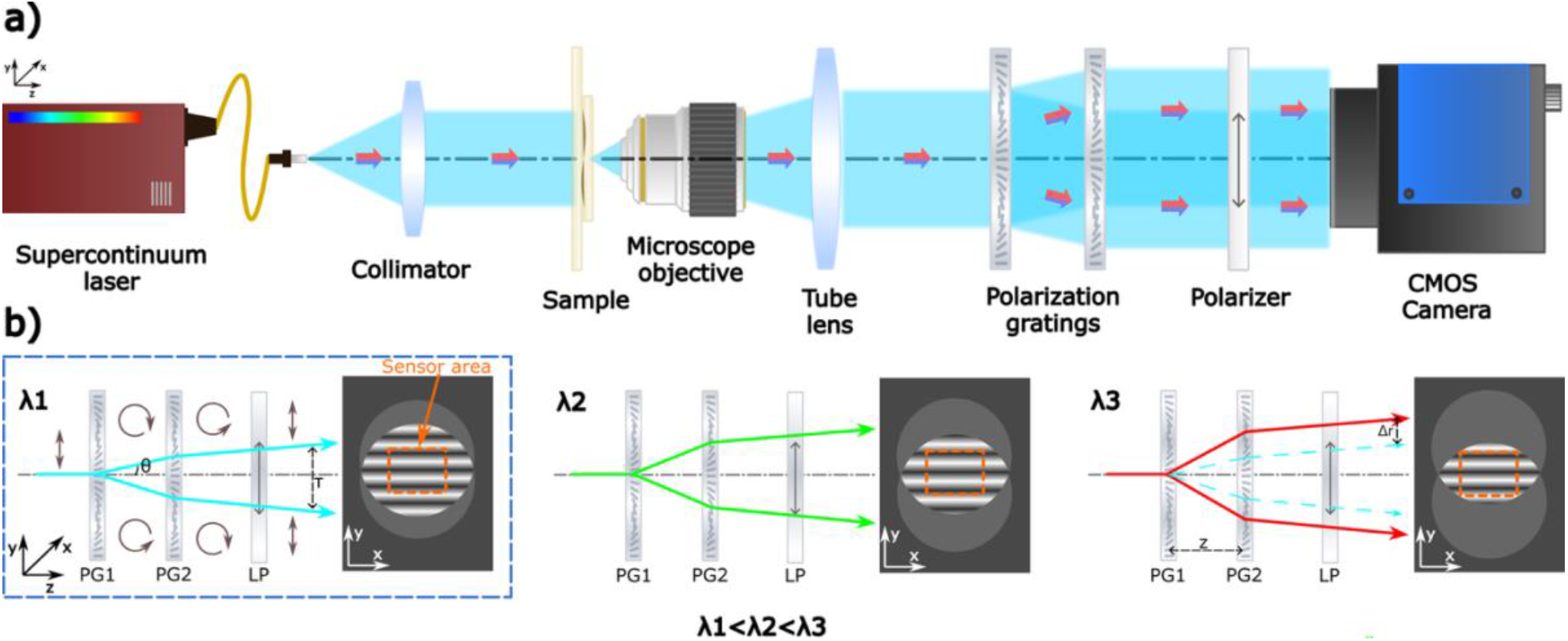
(a) Optical setup of a common-path holographic microscopy system based on polarization gratings for single-shot imaging. (b) Schematic of beam propagation through the grating module depending on the illumination wavelength *λ*. PG – polarization grating, LP – linear polarizer, *λ* – illumination wavelength, *τ* – shear between the beams, *θ* – diffraction angle, *z* – distance between the gratings, Δ*r* – beam displacement between frames. In the schematic marked with the blue frame, successive polarization states of the beam are schematically illustrated using brown arrows. For clarity, these markings are omitted in the remaining schematics; however, they apply to all cases.

The value of the shear *τ*_λ_ depends on the diffraction angle *θ*_λ_ and on the distance *z* between the gratings^46,53^ (Fig. 1(b). Variants of interferogram parameter modulation, i.e., translation and rotation of the gratings, are described in detail in the Methods section. When the shear value *τ* is larger than the camera sensor diagonal, the registered overlapping phase image will contain information from two disjoint areas of the object, effectively doubling the field of view. In order to recover undisturbed phase information from overlapped object replicas, it is necessary to record a series of at least *N* = 2 interferograms for different shear values *τ*.

In this work, we present a method for controlling the shear value *τ*_λ_based on modifying the diffraction angle *θ*_*λ*_. It depends, among other factors, on the wavelength of the incident light *λ*, via the diffraction grating equation *θ*_*λ*_ = sin^−1^(*λ*/*d*), where *d* is the grating period. The larger the wavelength *λ*, the larger the angle *θ*_*λ*_ and, consequently, the greater the shear *τ*.

Modulation of the shear value *τ*_*λ*_, by changing the wavelength *λ*, is implemented in two modes (Fig. 2):

**Fig. 2.**
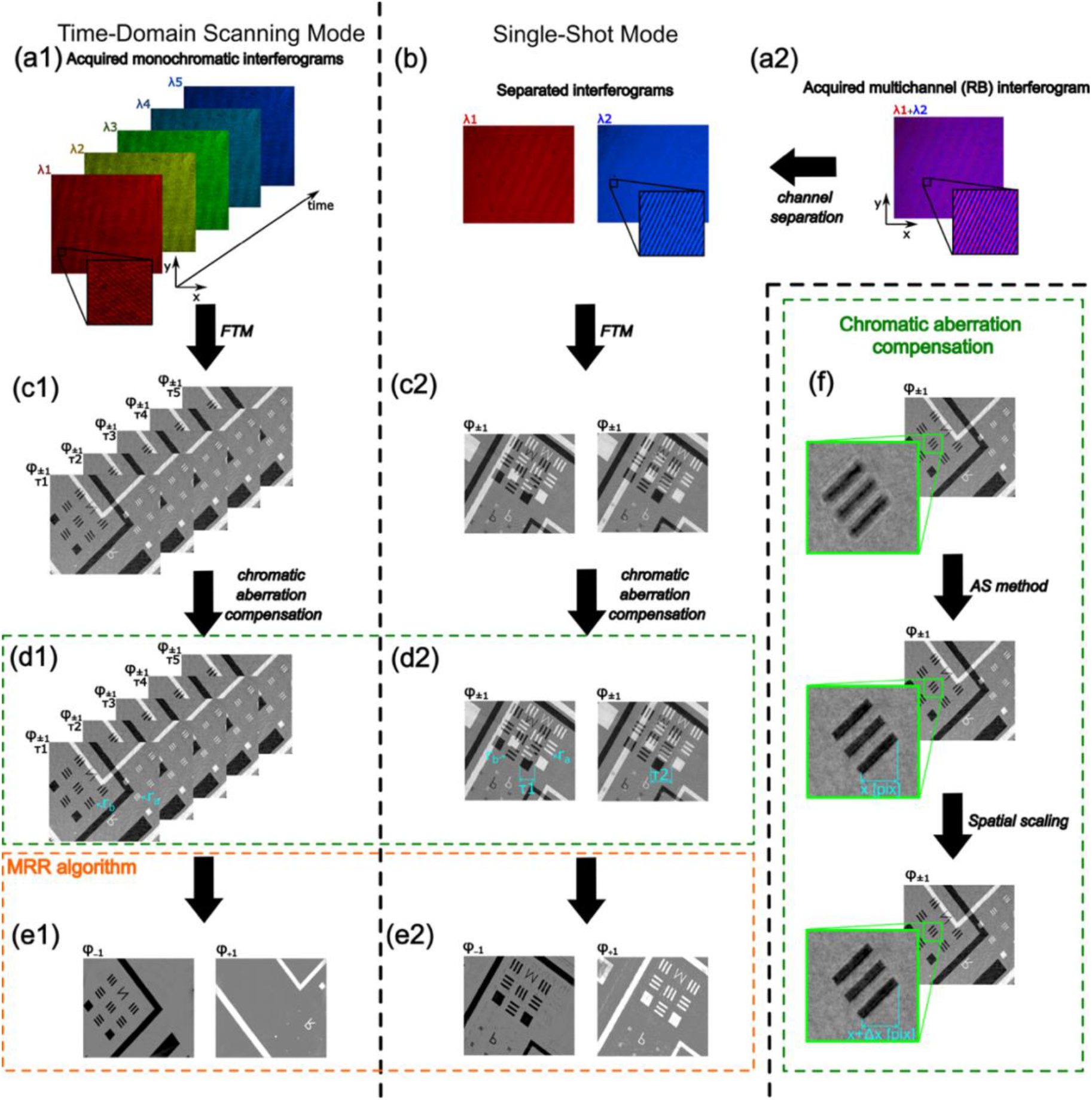
Data-processing workflow of the wsMRR algorithm for different measurement scenarios. (a1)–(e1) Processing of data acquired in the wavelength-scanning mode. (a2)–(e2) Processing of data acquired in the single-shot mode under dual-wavelength illumination. Panels (a) show the recorded interferograms with a zoomed-in view of the interference pattern. For the single-shot mode, (b) presents the set of interferometric images obtained by separating the color channels of the recorded frame. The next step involves reconstructing optical field using FTM: (c) display the phase information that can be recovered from this field. Sections (d) present the phase information *φ* ± 1 with visible replica overlap after correction of focus and magnification (see panel f), where *r*_*a*_ and *r*_*b*_indicate the phase encoded in the ±1orders, and *τ* represents the lateral spatial shift between the interfering beams. Finally, (e) show the separated phase reconstructions for both diffraction orders FOVs, obtained from a set of reconstructions corresponding to different values of the shift *τ*_*λ*_.

1. Wavelength scanning mode: Sequential illumination with different *λ* (step, e.g., Δ*λ* = 5 *nm*) and recording of the interferogram with a monochromatic camera after each change.

2. Single-shot mode: Illumination with two discrete *λ* (e.g., *λ*_1_ = 464 *nm* and *λ*_2_ = 630 *nm*) simultaneously. A single interferogram is recorded with a color (RGB) camera and then separated into the individual color channels. Only the red (R) and blue (B) channels are used for further processing, while the green (G) channel is omitted to limit the spectral cross-talk between the two wavelengths.

The application of wavelength scanning offers several advantages, such as higher stability and acquisition speed. However, it also requires accounting for dispersion resulting in chromatic aberrations (transverse and longitudinal) and necessitates additional processing steps.

Different wavelengths focus at slightly different planes (longitudinal chromatic aberration), introducing varying defocus between the recorded frames. Therefore, the reconstruction process begins with the application of the Fourier transform method to each frame in the series. The optical field recovered in this way is numerically refocused using the angular spectrum method^54^.

A change in wavelength also leads to a change in the system magnification (transverse chromatic aberration), resulting in a different mapping of the sizes of the same structures. This effect is corrected by applying geometric operations, consisting of image scaling using bicubic interpolation.

Subsequently, the phase value *φ*_±1_ recovered from each frame is scaled with respect to its corresponding wavelength. This scaling reflects the dependence of the phase shift on the wavenumber (*k* = 2*π*/*λ*) and represents a novel element introduced in the wsR2D-QPI method. Such a dataset, together with the estimated values of *Δr* - the difference in *τ* between frames - is processed using the MRR algorithm (described in detail in ^53^). This algorithm analytically separates the overlapping phase images (*φ*_+1_ and *φ*_−1_)from both replicas (Fig. 2).

### 2.2 Experimental results

#### 2.2.1 Comparison between R2D-QPI and wsR2D-QPI

To compare the impact of the replica shift variation method on imaging quality, we collected two datasets, each consisting of *N* = 20 interferograms recorded at different shear values.

The phase target (Lyncee Tec, Borofloat 33 glass with etched structures of 125 nm height) was imaged using a 5x/0.12 microscope objective and a low temporal coherence supercontinuum light source. In the first series (Figures. 3(a)-(d)), as a reference for the algorithm’s performance, we applied an R2D-QPI approach^53^. The change in the shear value *τ* was obtained by translating the diffraction grating (PG2) along the optical axis, with a step of *δ* = 0,5 *mm*, at a constant illumination wavelength *λ* = 535 *nm*.

**Fig. 3.**
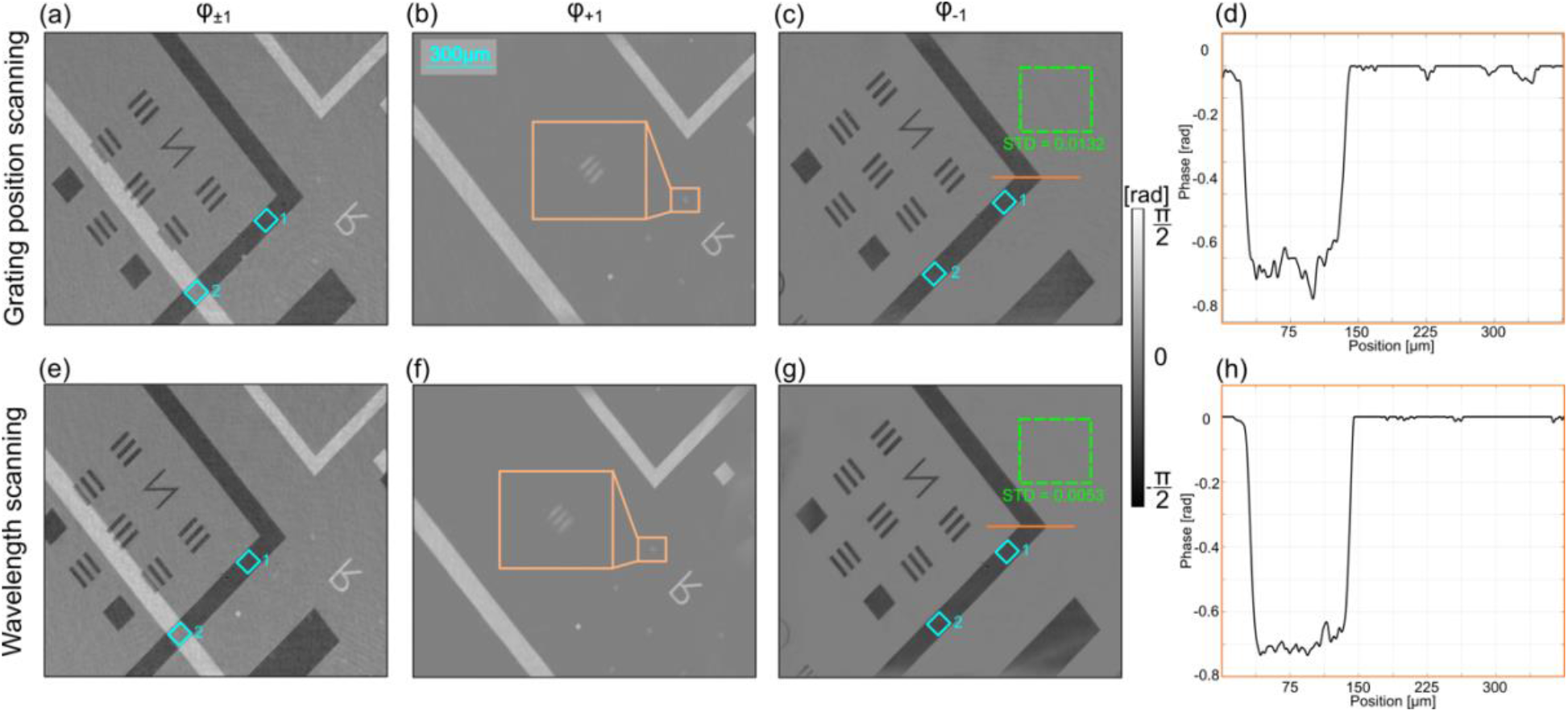
Comparison of phase-imaging results obtained for a phase resolution target using grid-shift scanning and wavelength-scanning approaches for a series of *N* = 20 frames. (a)–(d) Imaging results for varying replica displacement introduced by translation of the grating (a displacement between consecutive acquisitions *δ* = 0,5 *mm*). (e)–(h) Imaging results for varying replica displacement achieved through change of the illumination wavelength (the wavelength range 490 – 585 nm, sampled with a step Δ*λ* = 5 *nm*). (a), (e) Phase reconstructions *φ*_±1_ obtained using FTM, exhibiting visible overlap between the replicas. (b), (c), (f), (g) Reconstructed *φ*_+1_ and *φ*_−1_phase distributions, with the orange line indicating the cross-section shown in (d) and (h). Areas free of the object for which STD phase value was determined are indicated with green rectangles.

In the second series (Figures. 3(e)-(h)), we employed a proposed wsR2D-QPI in which the change in the shear value *τ*_*λ*_ was achieved by varying the illumination wavelength *λ*. The range of variation was from 490 nm to 585 nm with a spectral step of Δ*λ* = 5 *nm*. The adopted experimental parameters were selected to ensure a fair comparison between methods. Both the grating translation step in Series 1 and the spectral step in Series 2 generated a similar intra-replica shift value *Δr*, corresponding to Δ*r* ≈ 14 pixels.

Figures 3(a) and 3(e) present the initial phase maps obtained using the Fourier transform method. The recovered phase *φ*_±1_ corresponds to the phase difference *φ*_+1_ − *φ*_−1_ between two conjugated object replicas.

We performed the quantitative reconstruction of the separated phase distributions *φ*_+1_ and *φ*_−1_ (corresponding to the −1 and +1 diffraction orders) for the entire series (*N* = 20) using the MRR algorithm. Figures 3(b), (c), (f), and (g) present the results for both fields of view, covering the “N” and “R” groups, respectively.

The reconstructions obtained for both series are free from significant artifacts, which directly confirms that the new approach does not negatively affect imaging quality or the effectiveness of the phase retrieval process using the MRR algorithm.

To examine the influence of overlapping object replicas on the correctness of the phase reconstruction, the mean phase value and its standard deviation were calculated in two characteristic regions. The first of these is the region where fragments of the object overlap (Fig. 3, blue square no. 2), in which the phase information undergoes degradation as a result of superposition. The second is an area without overlap (Fig. 3, blue square no. 1), adopted as a reference point for correct phase representation. As indicated by the results summarized in Table 1, in both measurement scenarios the MRR algorithm enabled correct reconstruction of the phase value in the region with overlapping object replicas, despite the loss of information occurring there.

**Table 1.**
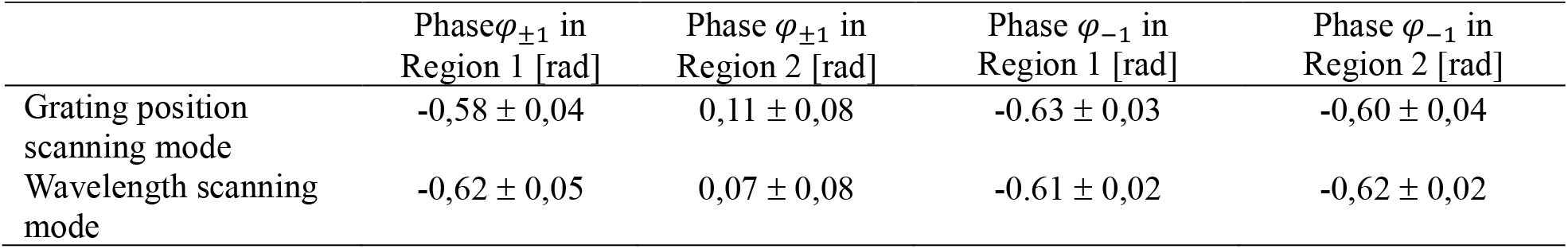
Average phase values along with their standard deviation for the areas marked with a blue rectangle in Fig. 5.

The key differences in quality between the modes become apparent during the analysis of the phase profile, marked with an orange line passing through one of the elements of the test structure in the reconstruction (Figures 3(c), (g)). The cross-sections shown in Figures 3(d) and (h), as well as the determined standard deviation (STD) of the phase values in object-free regions, clearly show that the reconstruction from Series 2 is characterized by a much more homogeneous background.

An analogous dependence was also observed under conditions of an elevated noise level. For this purpose, an additional experiment was carried out, imaging the same test object, but this time covered with a contaminated cover glass, which significantly degraded the signal-to-noise ratio. As indicated by the STD of phase values in object-free regions listed in Table 2, the application of the wavelength scanning mode enabled more effective averaging of these disturbances.

**Table 2.**
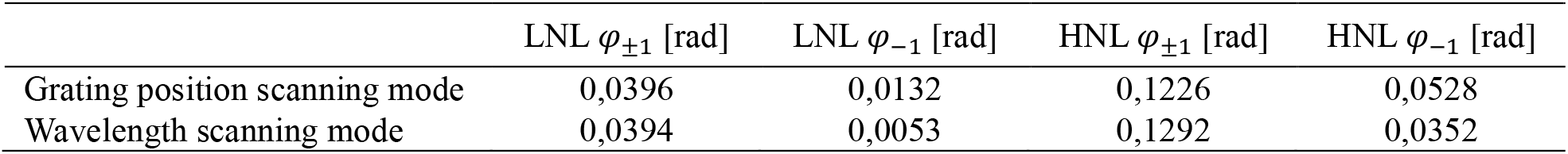
Phase standard deviation values of the single frame (*φ*_±1_) and the final reconstruction (*φ*_−1_) at low (LNL) and high noise levels (HNL)

#### 2.2.2 Parameter analysis in wsR2D-QPI

Within the experimental studies, the influence of key parameters (wavelength *λ*, spectral step *Δλ*, number of frames *N*, and total shift (*N* − 1) · *Δλ*) on the reconstruction quality was analysed. The obtained dependencies are presented in Figs. 4–7.

**Fig. 4.**
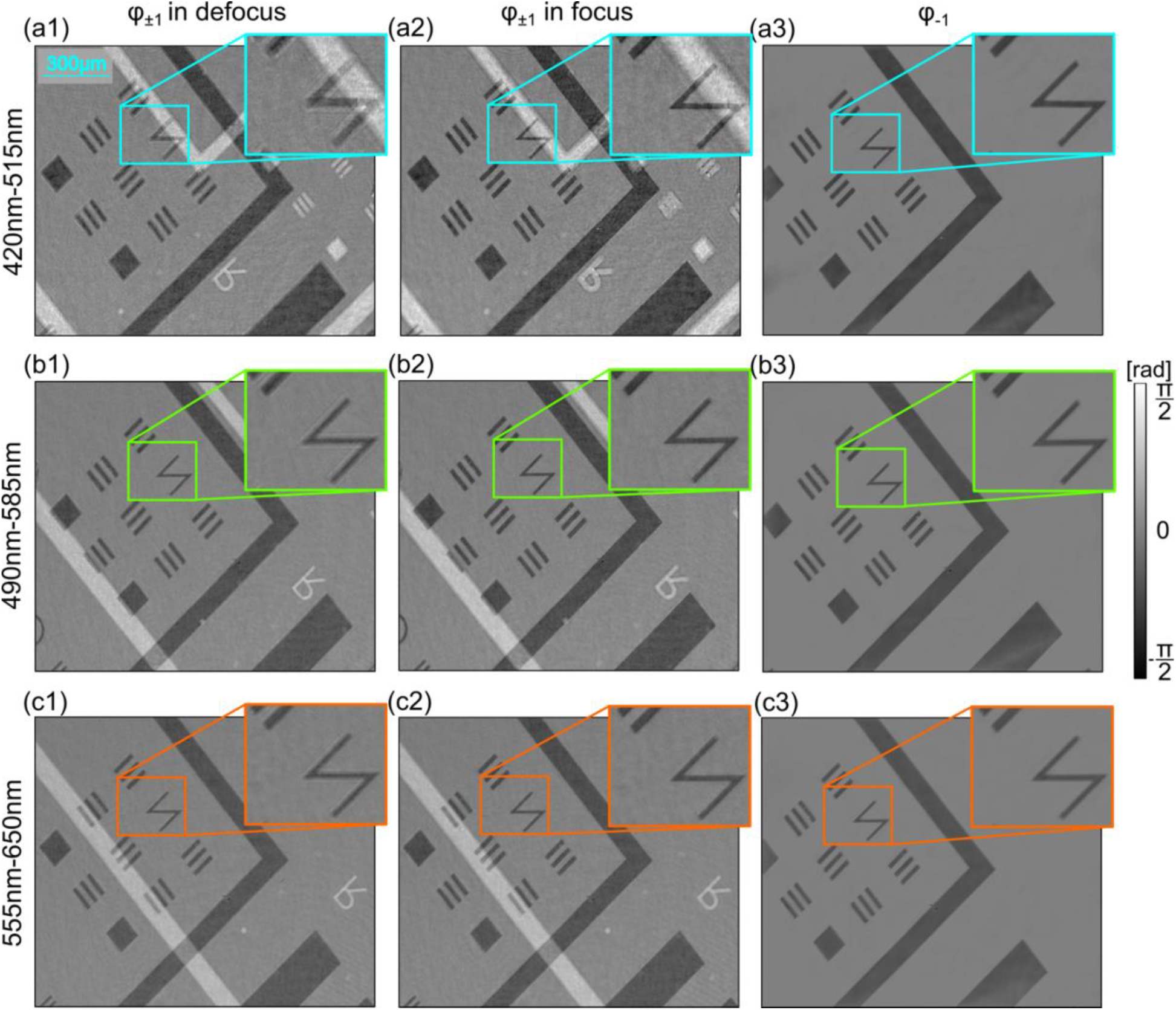
Comparison of phase-imaging results obtained for a phase resolution target from image series (*N* = 20) acquired over different illumination wavelength ranges. Phase reconstructions *φ*_±1_ retrieved using the FTM are shown for holograms before propagation to the object plane in (a1)–(c1) and for holograms after propagation to the focal plane in (a2)–(c2). The corresponding reconstructed *φ*_+1_ and *φ*_−1_ phase distributions are presented in (a3)–(c3). Results are shown for the following wavelength ranges: (a) 420–515 nm, (b) 490–585 nm, and (c) 555–650 nm.

In Figure 4, we present the results from three data series for different spectral ranges: 420– 515 nm (Figure 4(a)), 490–585 nm (Figure 4(b)), 555–650 nm (Figure 4(c)). Each series comprised *N* = 20 frames, recorded after changing the wavelength, with a step of Δ*λ* = 5 *nm* between consecutive frames. The focus of the imaging system was set for the central wavelength in each series.

The phase reconstructions *φ*_±1_ obtained exclusively from the first frame of each series, before propagation to the object plane, are shown in Figs. 4(a1)–4(c1), and after this propagation in Figs. 4(a2)–4(c2). Figs. 4(a3)–4(c3) present the final reconstructions of the phase *φ*_−1_. Despite the identical spectral spacing between the first and the central frame of each series, the object is most strongly defocused for the range with the shortest wavelengths (Figure 4(a1)). This is caused by the higher steepness of the objective dispersion curve in this range, which results in the same wavelength difference producing a larger change in the refractive index and, consequently, a more pronounced defocus effect. After numerical correction within proposed wsMRR framework (Fig. 4(a2)), this effect is compensated and the final phase *φ*_−1_ is reconstructed without defocus aberration (Fig. 4(a3)). This enables the use of shorter wavelengths, which guarantee better resolution, without the risk of image degradation due to dispersion, as its influence is corrected algorithmically.

Another parameter that significantly affects the reconstruction quality is the spectral step *Δλ*. Its value directly determines the magnitude of the beam shift *Δr* between consecutive frame. To illustrate the influence of this parameter on the reconstruction process, four measurement series were performed. Each series included the acquisition of two interferograms, spectrally separated by the specified values: 5 nm (Fig. 5(a)), 45 nm (Fig. 5(b)), 95 nm (Fig. 5(c)), and 160 nm (Fig. 5(d)). In each series, the wavelengths used to illuminate the sample were selected symmetrically with respect to the central wavelength equal to 545 nm.

**Fig. 5.**
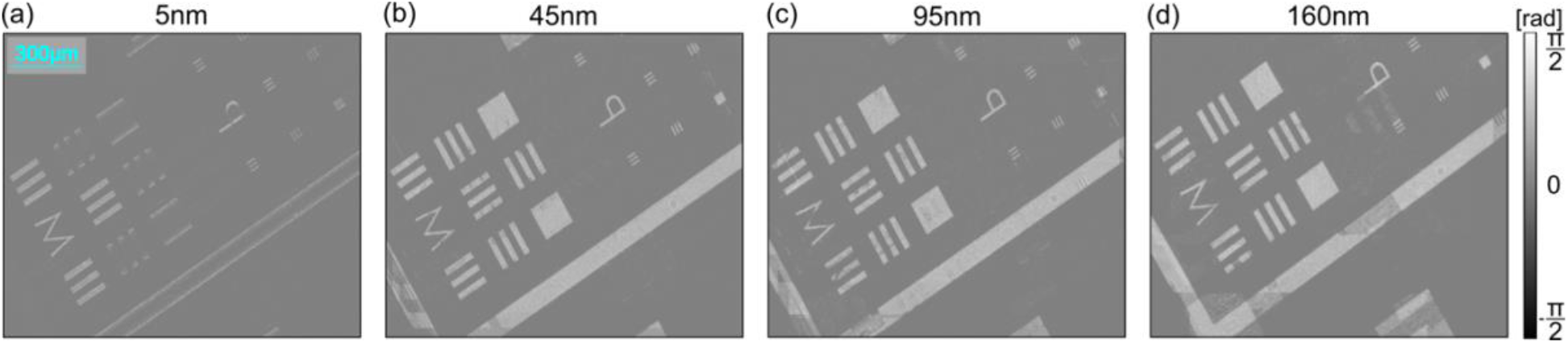
Comparison of phase-imaging results obtained for a phase resolution target from image series (*N* = 2) acquired with different spectral spacing between consecutive frames. Reconstructed *φ*_+1_ phase distributions for the +1 diffraction order FOV are shown for spectral spacings of (a) 5 nm, (b) 45 nm, (c) 95 nm, and (d) 160 nm between frames.

**Fig. 6.**
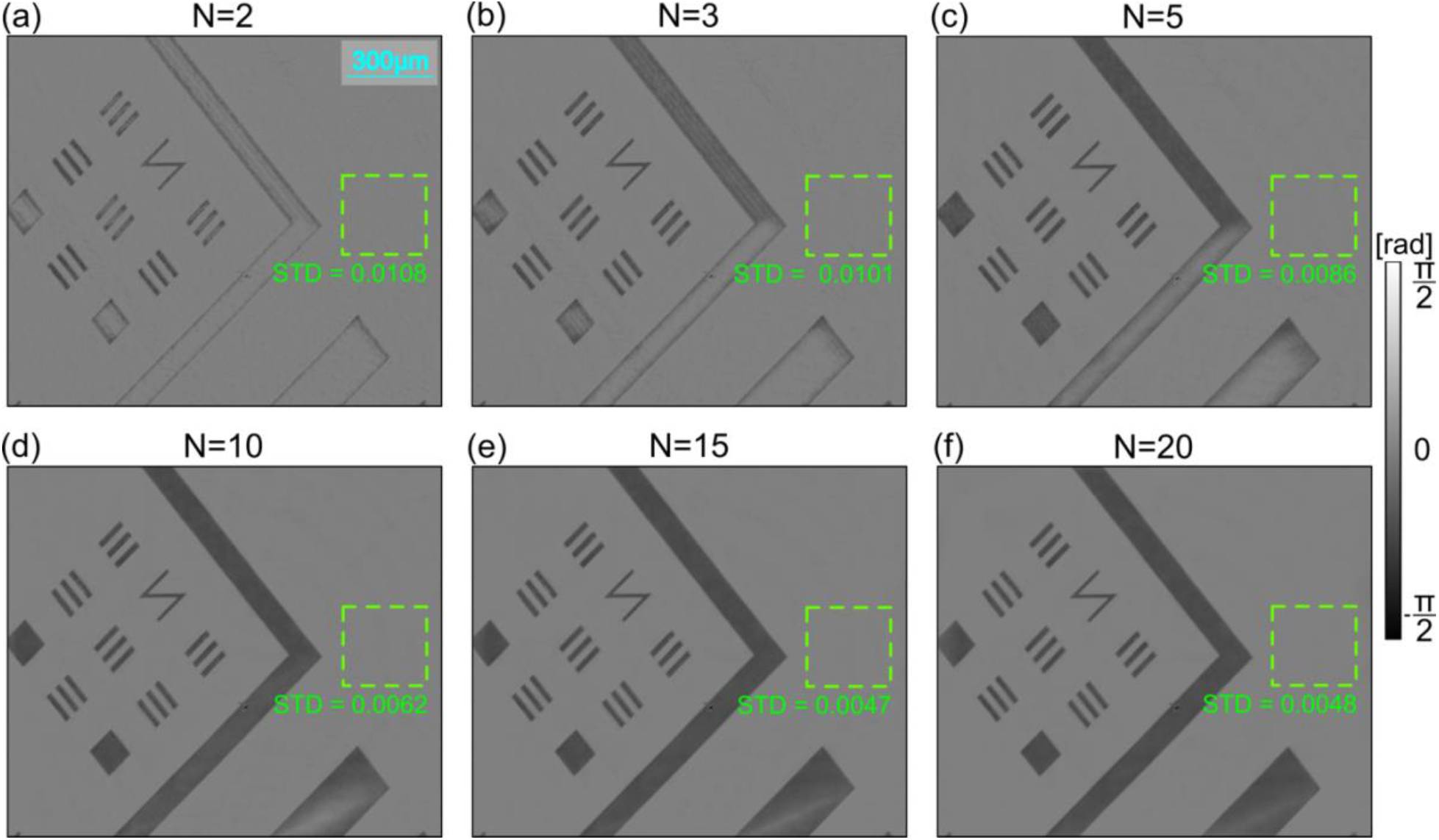
Comparison of phase-imaging results obtained for a phase resolution target from measurement series with varying numbers of frames acquired with an identical spectral step between frames (*Δλ* = 5 *nm*). Reconstructed *φ*_−1_ phase distributions for the −1 diffraction order fields of view are shown for series consisting of (a) *N* = 2, (b) *N* = 3, (c) *N* = 5, (d) *N* = 10, (e) *N* = 15, and (f) *N* = 20 frames. The STD phase value is calculated within the object-free region indicated by the green rectangle.

The phase reconstruction results *φ*_+1_ for the individual values of the spectral step *Δλ* are shown in Figure 5. We observed that an excessively large value of this parameter leads to a situation in which substantial parts of the object are visible only in a single frame from the entire measurement set. The consequence of this phenomenon is numerous artifacts in the reconstructed phase, particularly evident in the Figure 5(c) and (d). On the other hand, an excessively small value of the spectral step (Figure 5(a)) limits the algorithm’s ability to correctly reproduce the low-frequency components. In this case, only the edges of the object oriented perpendicular to the direction of the shift *Δr* are correctly reconstructed.

Another significant factor limiting the fidelity of low spatial frequency reconstruction is an insufficient number of recorded frames. The influence of this parameter on the reconstruction quality was investigated by performing measurement series with a constant spectral spacing *Δλ* = 5 *nm* between consecutive frames. The wavelengths were selected symmetrically with respect to the central value of 535 nm. Figure 5 shows the reconstructed phase image *φ*_−1_ of the replica for an increasing number of used interferograms: *N* = 2, 3, 5, 10, 15, and 20.

We observed that the smaller the number of frames, the more pronounced the difficulties in reproducing the low-frequency components, and additionally, the background inhomogeneity increases, as measured by the standard deviation. At the same time, as the number of frames increases, the improvement in reconstruction quality - in terms of low-frequency phase accuracy, resolution, and background uniformity - gradually saturates.

The progressively improved filling of large object areas observed with an increase in both the number of frames N and the spectral step *Δλ* is not accidental. The key importance lies in the value of their product, namely the total beam shift value defined as (*N* − 1) · *Δr*, rather than in the individual components of this parameter.

This relationship was confirmed by performing measurement series while maintaining a constant value of the total shift but with varying spectral sampling: *Δλ* = 90 *nm* (Figure 7(a)), 45 *nm* (Figure 7(b)), 30 *nm* (Figure 7(c)), 10 *nm* (Figure 7(d)), and 5 *nm* (Figure 7(e)). The obtained phase reconstruction results *φ*_−1_, shown in Figure 7, demonstrate that although the value of the spectral step decreases as the number of frames increases, the quality of low spatial frequency reproduction remains unchanged - precisely due to the constant value of the product (*N* − 1) · *Δr*.

**Fig. 7.**
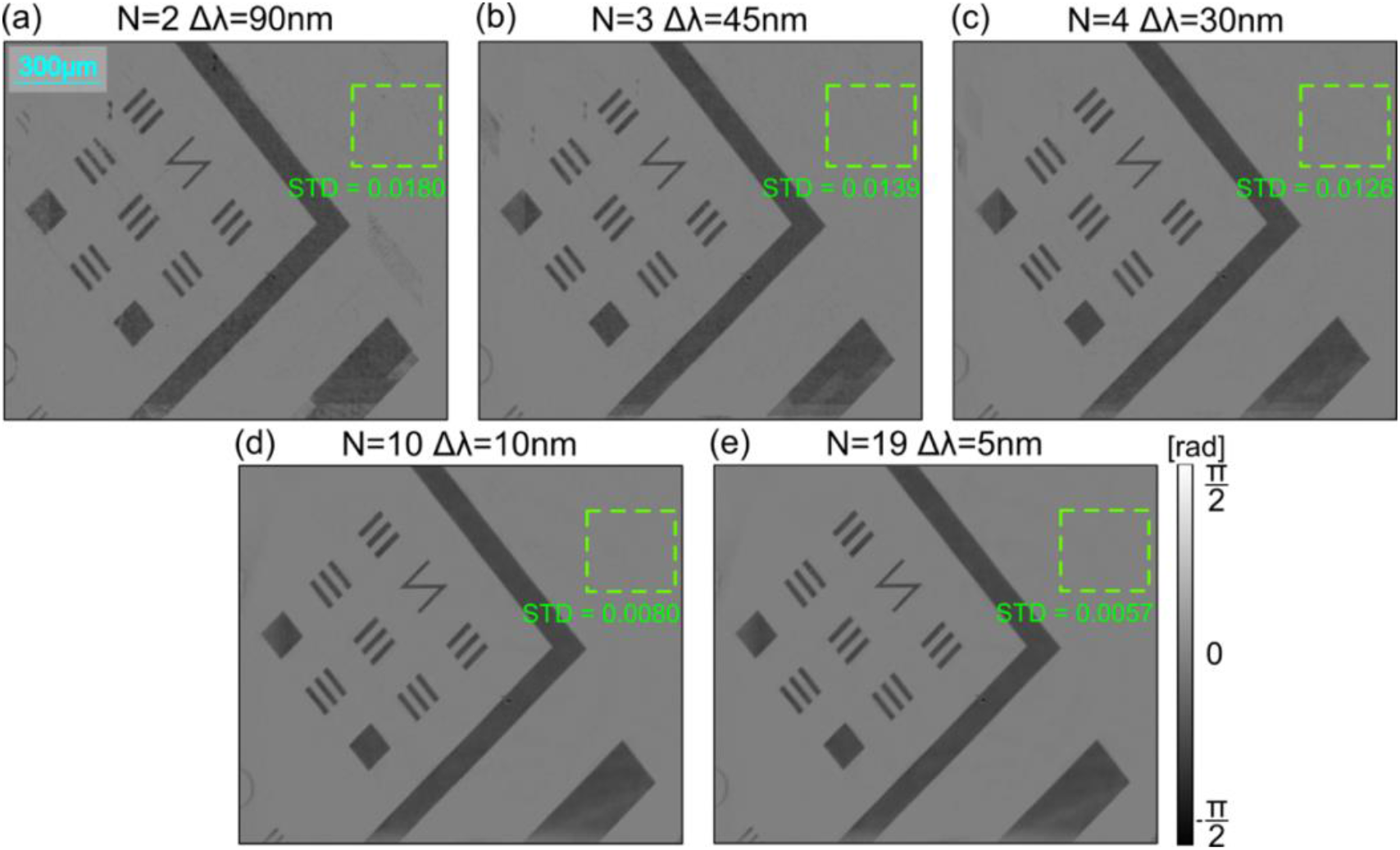
Comparison of phase-imaging results obtained for a phase resolution target from image series acquired with the same total shift value ((*N* − 1) · *Δr* = 90 *nm*) but with different sampling. Reconstructed *φ*_+1_ phase distributions for the +1 diffraction order FOV are shown for spectral spacings of (a) 90 nm, (b) 45 nm, (c) 30 nm, (d) 10 nm, and (e) 5 nm between frames.

In the case of a smaller number of frames and a large spectral step (Figure 7(a)), the reconstruction contains artifacts, caused by the visibility of substantial parts of the object only in a single frame (similarly as shown in Figure 5(d)). As shown by the calculated value of the STD in the background region, increasing the number of recorded frames provides an additional benefit in the form of noise averaging, which improves the signal-to-noise ratio in the resulting reconstruction. These observations lead to the conclusion that the acquisition of a larger number of frames with a smaller shift value *Δr* is generally a more advantageous solution. Nevertheless, the reproduction of the low-frequency components itself does not deteriorate relative to the remaining measurement series.

#### 2.2.3 Single-shot wsR2D-QPI

Next, we evaluated the method capabilities to remove the object replica using a single shot acquisition with color camera. Figure 8 shows the imaging results of technical and biological samples using the single-shot mode (Figs. 8(a1)-8(c2)). We present the phase reconstructions φ ±1 retrieved using the Fourier Transform Method (Figure 8(a)) together with the corresponding phase distributions (*φ*_+1_ and *φ*_−1_) for the ±1 diffraction order (Figure 8(b), (c)). Each measurement consisted of recording of a single frame, with the object illuminated by a beam composed of two wavelengths: 464 nm and 630 nm. Figs. 8(a1)-8(c1) present reconstructions of the phase target.

**Fig. 8.**
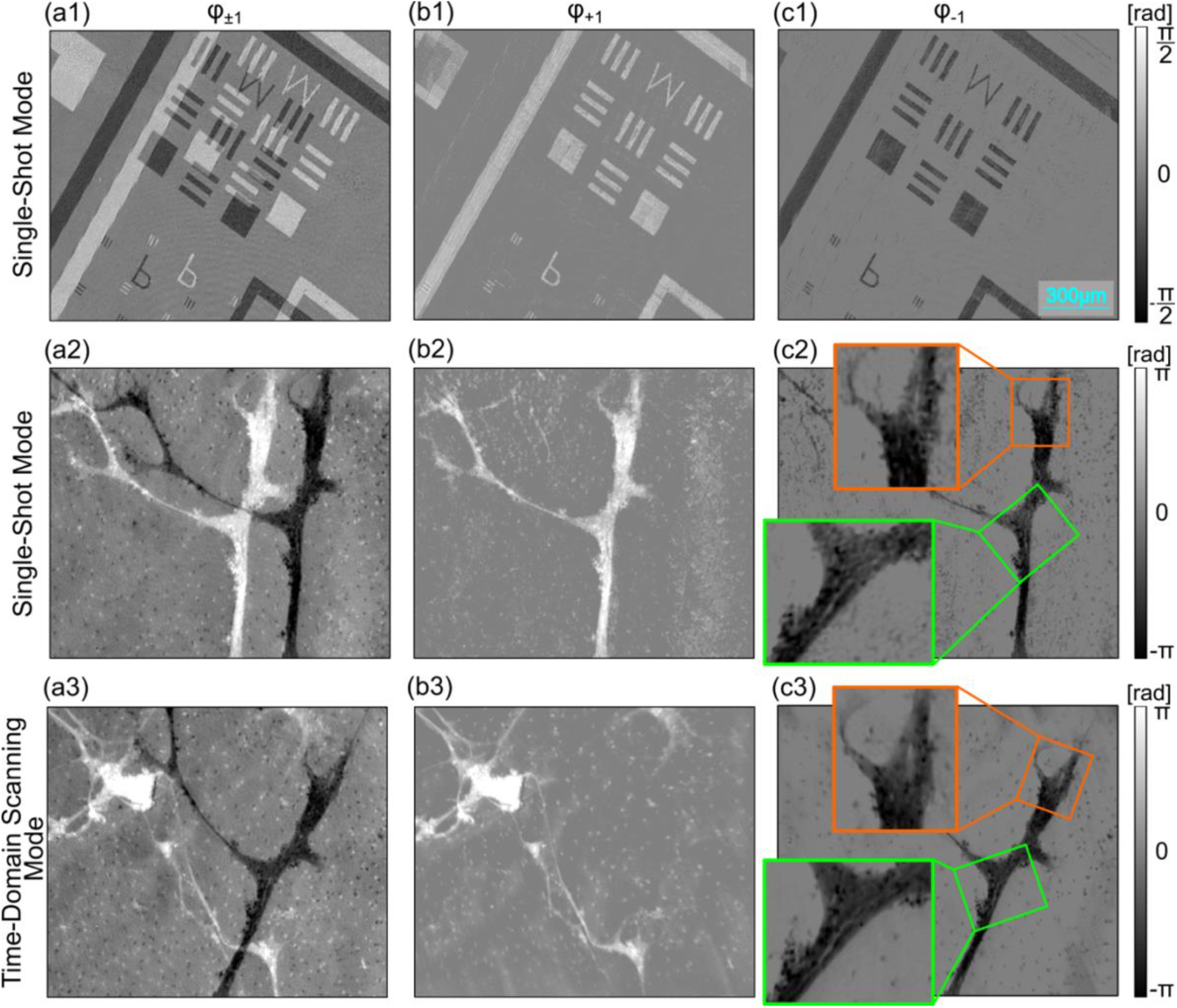
Single-shot phase imaging results compared to time-domain scanning mode results. Phase reconstructions *φ*_±1_ retrieved using the FTM with visible replica overlap, along with the corresponding phase distributions (*φ*_+1_ and *φ*_−1_) for the ±1 diffraction order FOV, are presented for a phase resolution target in (a1)–(c1) and for a biological sample (neuronal cell culture) in (a2)–(c3). Measurements were performed in single-shot mode using illumination wavelengths of 464 nm and 630 nm in (a1)–(c2), and in time-domain scanning mode across a wavelength range of 470–622 nm (sampled with an Δ*λ* = 8 *nm* step) in (a3)–(c3).

In the reconstructions, no artefacts arising from the limited number of frames (*N* = 2) at a large spectral step were observed; such artefacts were visible in the corresponding measurement series acquired in the wavelength-scan mode (Fig. 5(d)). They were mitigated by reducing the grating rotation angle *β*, since it was not possible to change the spectral step (due to the use of a camera with a Bayer filter) or increase the number of frames (because of cross-talk from the G channel). At a smaller angle *β*, for the same change in wavelength, the displacement *Δr* between frames is smaller, meaning that a smaller portion of the object appears in only one field of view. Consequently, the reconstruction contains fewer artefacts; however, this limitation reduces the carrier frequency and thus worsens the separation of the signal from the direct-current (DC) component in the spectrum, which may negatively affect phase reconstruction.

In the single-shot mode, we additionally imaged a biological object in the form of a neuronal cell culture 8(a2)-8(c2). Both the axons themselves and the small cells in the background, which only weakly modulate the phase, were reproduced correctly. For comparison, we imaged the same object using the wavelength scanning method 8(a3)-8(c3), recording *N* = 20 frames within a similar spectral range of 470 - 622 nm (step Δ*λ* = 8 *nm*). It should be emphasized that the measurement parameters for both modes were independently optimized to ensure the best possible imaging quality in each of them. This provides a basis for a reliable comparison of the maximum achievable imaging quality in each of these measurement configurations. Due to the necessity of changing the grating rotation angle *β*, the phase reconstructions *φ*_+1_ from the two modes represent different fields of view (Fig. 8(b2),8(b3)). In contrast, the phase reconstructions *φ*_−1_ cover the same field of view (Fig. 8(c2),8(c3)), which enables a comparison of the quality of both methods. The performed analysis indicates that although the imaging quality in the single-shot mode is lower, most details and fine structures are visible in both reconstructions.

It should be emphasized, however, that the wavelength scanning mode, owing to the ability to freely shape the measurement parameters - including the key number of frames - enables the achievement of significantly higher reconstruction quality for the examined objects.

Despite the slightly lower imaging quality compared to the wavelength-scanning method, the undeniable advantage of the single-shot mode remains the significantly better temporal resolution and the ability to observe dynamic objects. To demonstrate this capability, we examined a sample containing moving yeast cells (Figure 9). The sample was illuminated simultaneously with two wavelengths - 464 nm and 630 nm - selected due to the high sensitivity of the color camera in these spectral ranges and imaged using a 40×/0.65 microscope objective.

**Fig. 9.**
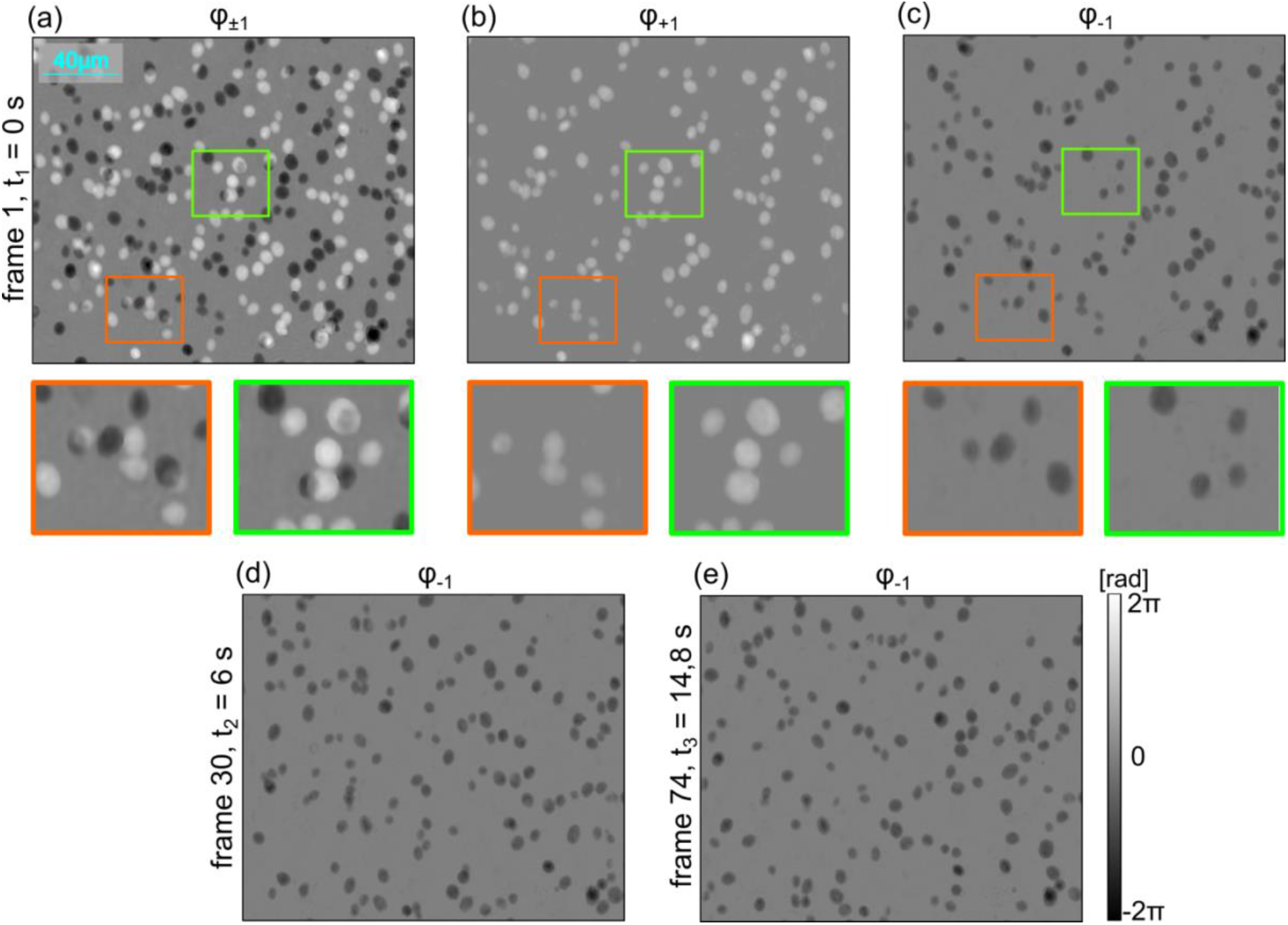
Single-shot phase-imaging results of a dynamic object (yeast cells). (a) Phase reconstructions *φ*_±1_ retrieved using the FTM. (b-e) Corresponding phase distributions (*φ*_+1_ and *φ*_−1_) for the ±1 diffraction order fields of view: (b,c) first frame in the recorded sequence, (d) 30th frame, and (e) 74th frame.

Figure 9(a) shows the phase reconstruction with the fields of view from both beams superimposed. In the magnified regions marked in green and orange, it is evident that the high density of the sample leads to local replica overlap in the phase reconstruction. Nevertheless, as shown by the separated phase reconstructions *φ*_+1_ and *φ*_−1_ (Figs. 9(b) and 9(c)), the algorithm readily recovers both fields of view - without loss of information or the introduction of additional artifacts.

Figure 9(c) shows the first recorded frame (t = 0 s), whereas Figures 9(d) and 9(e) present the 30th (t = 6 s) and 74th (t = 14.8 s) frames of the recorded sequence, respectively (frames acquired every 0.2 s).

## Discussion

In this work, we presented the wsR2D-QPI – a new approach enabling the separation of overlapping fields of view in common-path DHM systems– that. The essence of the method is the application of the wsMRR algorithm to a series of holograms recorded with different shear between the interfering beams. This makes it possible not only to retrieve the phase but, above all, to separate the object information from its replicas, which effectively results in the extended field of view.

The key innovation is the introduction of control over the shear value through changes in the wavelength of light, allowing the singl-shot replica free QPI, crucial for the imaging of dynamic processes. In contrast to previous solutions based on mechanical scanning^50–53^, the new approach simplifies the system architecture, increases the stability and repeatability of measurements, and enables more effective noise averaging. However, changing the wavelength introduces additional effects related to chromatic aberrations. To mitigate them, additional hologram processing stages were introduced to minimize the influence of these distortions on the reconstructed phase. An additional advantage of the proposed method is the possibility of creating a hologram series also by splitting a single frame into color channels. This makes it possible to obtain the dataset required for phase reconstruction in a single acquisition. Moreover, unlike the CIM method, which uses high-pass filtering^50–52^, this technique does not require such an operation. As a result, the low-frequency components of the phase are better preserved, which promotes a more faithful representation of the investigated objects.

Two measurement modes were developed that optimize the measurement procedure for different applications. The wavelength time-domain scanning mode allows flexible adjustment of the acquisition parameters (spectral step, number of frames) to the characteristics of the sample. This translates into higher imaging quality and good representation of the low-frequency phase components. In turn, the single-shot mode offers excellent temporal resolution, enabling the observation of dynamic processes while still maintaining high fidelity of phase reconstruction.

However, in order to minimize reconstruction artifacts, it is necessary in single-shot mode to reduce the carrier frequency. From the perspective of the FTM technique, this is unfavorable because the information peak is less well separated from the direct current (DC) component, which may potentially reduce the quality of the reconstructed phase.

Another aspect requiring attention, and opening the way for further improvement of the method, is the implementation of color-channel crosstalk correction^55^. This would allow the effective use of three wavelengths (including the eliminated green channel), which would increase the number of frames recorded in a single acquisition in the single-shot mode and thereby improve the quality of phase reconstruction.

In summary, the presented wsR2D-QPI approach constitutes a versatile tool for quantitative phase imaging in common-path shearing configurations. It enables analytical and complete removal of unwanted contributions from replicas, even when imaging dense samples. Thanks to two operation modes, it can be applied both to imaging static biological samples, enabling detailed representation of their structure (e.g., tissue architecture), and to observing rapidly changing processes in cell cultures, where excellent temporal resolution is crucial. The proposed method constitutes a promising tool for a wide spectrum of biomedical applications.

## Methods

### 4.1 Setup details

In the presented setup, the light source is a partially coherent, tunable laser (supercontinuum laser, NKT Photonics SuperK EVO HP EU-4, FWHM ≈ 5 nm), enabling sequential wavelength scanning or simultaneous illumination with multiple wavelengths. The beam passes through the sample, the objective (Zeiss 5x/0.12 or Nikon 40x/0.65), the tube lens, and reaches the polarization grating assembly (period of the PG *d* = 6.3 *μm*).

The polarization state of the generated diffraction orders depends on the polarization of the incident beam. Under linearly polarized illumination, two diffraction orders (±1) are produced with orthogonal circular polarizations (Fig. 1b). In the case of circular polarization, the grating converts it into the orthogonal one^56^. Two identical gratings arranged in series introduce a transverse beam shift *τ*.

Its value depends both on the diffraction angles *θ* at the two gratings and on the distance *z* between them. The first grating (PG1) splits the incident beam, transforming its polarization into circular polarization and deflecting the individual diffraction orders at angles ±*θ*_*λ*1_. According to the grating equation, the diffraction angle *θ*_*λ*_ depends on the wavelength *λ* and the grating period d. Consequently, the longer the wavelength, the larger the diffraction angle *θ*_*λ*_ and, therefore, the greater the mutual separation of the beams *τ*_*λ*_.

The second grating (PG2) compensates for this deflection; however, the angles ∓*θ*_*λ*2_ at which it diffracts the beam also depend on the mutual rotation angle between the line of the two gratings *β*. When the gratings are not aligned in parallel (*β* ≠ 0), the beams propagate relative to each other at an angle (*θ*_*λ*1_ − *θ*_*λ*2_). This phenomenon enables precise control of the fringe spatial frequency in the interferometric image through mutual rotation of the gratings. In the measurements, the introduced carrier frequency had a value typical for the FTM method.

The beam separation *τ* is also influenced by the distance *z* between the gratings. Reducing this distance shortens the propagation path of the beams at an angle, which leads to a smaller wavefront displacement *τ*. To enable interference of orthogonally polarized beams, a linear polarizer (LP) is placed in front of the camera. The interference pattern is recorded by a CMOS sensor.

To separate the undisturbed phase information from both replicas (orders), it is necessary to record a series of at least *N* = 2 interferograms for different values of *τ*_*λ*_. This is implemented in two modes: (i) wavelength scanning and (ii) single-shot.

The first mode consists of sequential illumination of the sample with different wavelengths *λ* and recording interferograms with a monochrome camera (Daheng Mer2-301-125U3M-L, resolution 2048×1536, pixel size 3,45 µm) after each change. The number of interferograms in the series, the spectral range, and the spectral step are selected depending on the requirements. In the second mode, the sample is illuminated simultaneously with discrete wavelengths *λ* = 464 *nm* and *λ* = 630 *nm*. A color (RGB) camera (Daheng Mer2-301-125U3C-L, resolution 2048×1536, pixel size 3,45 µm) records a single interferogram, which is separated into color channels. Due to cross-talk in the green channel, only data from the B and R channels, corresponding to the applied wavelengths, are used for further reconstruction.

As a reference method for changing the replica shear, the approach described in the previous work^53^ was used. In this variant, the change in shear value *τ* results from changing the distance between the gratings. During the experiment, one of the gratings was precisely translated with a step *δ* = 0,5 *mm* along the optical axis using a motorized linear stage (ThorLabs MTS25/M-Z8). In this way, at a constant illumination wavelength *λ* = 535 *nm*, a series of *N* = 20 interferograms was recorded using a monochrome camera (Daheng Mer2-301-125U3M-L, resolution 2048×1536, pixel size 3,45 µm).

### 4.2 Data Processing Pipeline and Reconstruction

The reconstruction process is based on a procedure for retrieving and analytically separating overlapping phase images originating from both beam replicas. For this purpose, the MRR algorithm^53^ is used in a new implementation, wsMRR, extended with a preprocessing stage compensating for dispersion effects. The complete data-processing pipeline is schematically presented in Figure 2.

The reconstruction process begins with the application of the FTM to each recorded interferogram in order to recover the optical field. Then, using the Angular Spectrum Method (ASM)^54^, the field is numerically propagated to the objective focal plane. This step is necessary because the system focus changes with the illumination wavelength, while the sample position remains unchanged during the measurement series, which results in a defocused image for some wavelengths. In addition, changes in wavelength cause magnification variations, which are corrected through geometric image transformations based on characteristic sample features. The obtained phase maps are also scaled with respect to the wavelength *λ*, due to the dependence of the phase value on this parameter.

From the optical field prepared in this way for each frame, the phase *φ*_±1_ corresponding to the phase difference between the two interfering replicas, *φ*_+1_ and *φ*_−1_, is retrieved. These values, together with the displacement *Δr* (the difference in *τ*_*λ*_ between frames) estimated from sample features, constitute the input data for the main part of the MRR algorithm.

In the first step, the algorithm estimates the individual phases 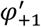 and 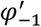 by compensating for shifts and averaging the obtained dataset 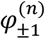. Then, in a two-stage procedure, based on the estimates 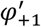 and 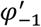 from opposite orders, the algorithm estimates artifacts *ξ*_+1_ and *ξ*_−1_ in the form of blurred replicas along the *Δr* direction.

The phases reconstructed in this way, 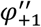 and 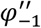, are analytically cleaned of the image of the opposite replica but still contain a component *ψ*_±1_ originating from the same diffraction order. To recover fully quantitative phase information, an iterative correction is applied, which progressively filters out the unwanted *ψ* component. A detailed description of the MRR algorithm implementation is presented in ^53^.

## Acknowledgements

This work was funded by the National Science Center Poland (2024/55/D/ST7/02792). The study was conducted on devices co-funded by the Warsaw University of Technology within the Excellence Initiative: Research University (IDUB) program.

## Author Contributions

A.P.: Investigation, Visualization, Writing – original draft. M.R.: Methodology, Software, Data curation, Formal analysis, Validation, Writing – review & editing. M.S.: Resources. M.T.: Supervision, Conceptualization, Resources, Writing – review & editing. P.Z: Methodology, Supervision, Conceptualization, Resources, Funding acquisition, Writing – original draft, Writing – review & editing.

## Supplementary information

Supplementary information for this paper is available at https://doi.org/10.29026/xxx.20xx.xxxxxx

